# TLS-Sim: An Open-Source Toolkit for Individualized Transcranial Light Stimulation Modeling with Deep-Learning-Accelerated Simulation

**DOI:** 10.64898/2026.09.09.749356

**Authors:** Keyao Zhang, Hai Jia, Zhilin Li, Shaodi Wang, Yushuai Zhang, Fengyu Cong, Zaixu Cui, Xiaoli Li, Chenguang Zhao

## Abstract

**Background:** Accurate modeling of photon transport through the heterogeneous tissues of the human head is important for individualized transcranial light stimulation (tLS). However, high-precision Monte Carlo (MC) simulations can be computationally demanding, and existing general-purpose simulation tools do not provide an integrated, brain-oriented workflow.

**Methods:** We developed TLS-Sim, a framework that integrates subject-specific head-model construction, multiple light-source configuration, GPU-accelerated MC photon simulation, visualization, and intracranial dosimetric analysis. Its central feature is a deep-learning-accelerated MC pathway, termed Flux-to-Flux, that estimates a high-photon-count energy-deposition field from a paired low-photon-count simulation. Sparse and full Monte Carlo fields were encoded using a three-dimensional variational autoencoder (VAE), and a three-dimensional velocity network was trained to transform the sparse latent representation toward the corresponding full-simulation representation.

**Results:** We demonstrate the end-to-end toolkit and the Flux-to-Flux (F2F) pathway on a representative individualized head model evaluated across three source configurations. Relative to the sparse simulation, the learned prediction improved agreement with the full Monte Carlo reference by approximately +4 dB in signal-to-noise ratio on average, with correspondingly higher structural similarity and lower mean absolute error.

**Conclusions:** TLS-Sim provides a modular, openly released computational framework for individualized tLS simulation and analysis. The Flux-to-Flux pathway is presented as an implemented capability that reduces the computational burden of high-photon-count simulation while improving spatial agreement with the full MC reference, most strongly at the low-fluence field boundaries where the sparse simulation is noisiest.

## 1. Introduction

Transcranial light stimulation (tLS), commonly framed as transcranial photobiomodulation (tPBM), delivers red or near-infrared light through the scalp and skull to modulate biological processes within the brain [1–3]. Human studies have demonstrated beneficial effects of tLS across multiple domains, including attention, executive function, memory, and emotional processing, together with increased cerebral oxygenation and cytochrome-c-oxidase activity [4–9]. The intracranial distribution of photon energy deposition is jointly determined by the wavelength, source geometry, source position and orientation, tissue optical properties, and subject-specific anatomical characteristics [10–12]. Consequently, stimulation protocols with the same parameters may produce different patterns of intracranial photon deposition across individuals, potentially affecting both stimulation efficacy and safety [10, 13, 14]. Computational modeling is therefore essential for estimating energy delivery to target regions, comparing stimulation protocols, and deriving physically interpretable dosimetric measures [14, 15].

Monte Carlo methods are widely used to model photon transport in heterogeneous biological media [11, 16–19]. General-purpose Monte Carlo simulators can accommodate complex anatomical geometries, including body models; however, they are not specifically designed to address the complete modeling requirements of transcranial light stimulation. Accurate tLS dosimetry requires the coordinated representation of subject-specific anatomy, wavelength-dependent tissue optical properties, source position and orientation, stimulation parameters, and high-photon-count energy-deposition field. In practice, these procedures are often distributed across multiple software tools, increasing the technical burden and making individualized dose estimation, protocol comparison, and reproducible implementation difficult.

Fast field estimation is a prerequisite for iterative source-position optimization and interactive target navigation, which are impractical with repeated full Monte Carlo simulation. Studies in transcranial magnetic and direct-current stimulation have pursued this goal by using deep learning to approximate subject-specific electric-field or current-density distributions in near-real-time [20, 21], demonstrating that the cost of an expensive physical simulation can be partially shifted to an offline training stage. In the tLS domain, GPU-accelerated platforms such as Monte Carlo eXtreme (MCX) substantially reduce the computational time required for three-dimensional photon-transport simulations [14, 17]. Nevertheless, high-photon-count simulations remain burdensome [22, 23] when repeated across subjects, source locations, incident directions, wavelengths, or optimization iterations. This limitation is particularly important for individualized tLS planning, in which numerous candidate stimulation configurations may need to be evaluated.

A network that maps anatomy directly to the photon field would have to learn tissue segmentation, optical-property assignment, and source geometry all at once. This entangled cross-modal problem is hard to train and tends to fail when applied to new anatomy. A more robust alternative is to generate two corresponding optical fields [24]: an approximate field from a low-photon-count Monte Carlo simulation and a low-noise, high-fidelity reference field from a high-photon-count simulation. The network is then trained to learn the mapping from the approximate field to the reference field, forming a F2F task in which the input and output represent the same physical quantity. Because this physical prior is already contained in the input, training is more stable and the model extrapolates more reliably—an advantage for the demanding multilayer scalp-skull-CSF-cortex light path of tLS.

We developed TLS-Sim as a modular framework for individualized tLS simulation. Within this framework, we implemented the F2F strategy described above as a concrete pathway: a fast low-photon-count (sparse) Monte Carlo field serves as the input, and the network is trained to predict the corresponding high-photon-count (full) field. The sparse and full fields describe the same physical quantity, and the sparse field already carries the subject’s anatomy, tissue properties, and source configuration. The learning task is therefore to restore a clean, high-fidelity field from a noisy, under-sampled one, rather than to generate a field from anatomy.

This study had two objectives: first, to describe the design and implementation of TLS-Sim as an open-source toolkit; and second, to demonstrate the F2F pathway on a representative individualized head model and to evaluate whether the reconstructed field showed improved agreement with the high-photon-count Monte Carlo reference relative to the original low-photon-count simulation, while reducing the computational cost of field estimation.

## 2. Materials and Methods

### 2.1 Study design and TLS-Sim overview

TLS-Sim was developed as an open-source, modular Python framework for individualized modeling and dosimetric analysis of transcranial light stimulation. Its workflow consists of four principal stages: (i) construction of a voxel-based anatomical head model, (ii) specification of light-source geometry and tissue optical properties, (iii) simulation of intracranial photon energy deposition, and (iv) visualization and quantitative analysis of the resulting energy-deposition field.

TLS-Sim provides two computational pathways for photon-transport simulation. The conventional pathway performs GPU-accelerated Monte Carlo simulation. The accelerated pathway employs a F2F model to reconstruct a high-photon-count energy-deposition field from a computationally inexpensive low-photon-count simulation. The low- and high-photon-count fields are generated under identical anatomical, optical, and source conditions, enabling the model to suppress sampling noise while preserving the underlying spatial pattern of photon propagation.

Individual modules can be executed sequentially as an end-to-end workflow or independently for specific tasks. This modular design provides a flexible foundation for subsequent source optimization and target-navigation applications.

### 2.2 TLS-Sim software implementation and graphical user interface

TLS-Sim was implemented in Python using a modular software architecture that integrates functional components for anatomical preprocessing, optical simulation, F2F reconstruction, dosimetric analysis, and visualization within a unified workflow. SimpleITK [25], nibabel [26], and SciPy [27] were used for volumetric image input/output, resampling, coordinate transformation, and numerical image processing. NumPy [28] supported multidimensional array operations and numerical computation, whereas pandas [29] was used for organizing, processing, and exporting tabular simulation and dosimetric results. SuperSynth [30, 31] and SynthMorph [32–37] supported tissue segmentation and spatial registration, respectively, whereas pmcx [17] provided the Python interface to MCX for GPU-accelerated Monte Carlo photon-transport simulation. The F2F model was implemented using PyTorch [38] and MONAI [39]. Matplotlib [40] and PyVista [41] were used for two-dimensional and interactive three-dimensional visualization.

TLS-Sim was operated through a graphical user interface implemented using PyQt6 (Riverbank Computing). The interface was organized into dedicated workspaces for anatomical-image preprocessing, optical simulation, voxel-level field inspection, and depth-dependent attenuation analysis. Users could select or generate anatomical head models, define light-source geometry and position, assign tissue optical properties, specify photon counts and simulation parameters, execute Monte Carlo simulations or F2F reconstruction, and inspect the resulting energy-deposition fields without directly editing MCX configuration files or Python scripts. The graphical interface provided access to the underlying computational modules without requiring users to directly modify backend configuration files.

Volumetric data were exchanged between processing modules using NIfTI files, while simulation settings and processing information were stored in structured configuration and metadata files. Anatomical models, segmentation results, simulation outputs, and derived analysis results were organized in dedicated processing directories. Relevant simulation parameters and processing information could be exported through the reporting function to facilitate retrospective inspection and reproducibility.

### 2.3 Anatomical data and individualized head-model construction

TLS-Sim supports both built-in anatomical templates and subject-specific head models derived from structural T1-weighted magnetic resonance imaging data. The built-in resources include the ICBM 152 Nonlinear Asymmetrical template (MNI152NLin2009cAsym) [42, 43], the MNI-Colin27 anatomical model [44], and five age-specific MNI Pediatric Asymmetric templates spanning approximately 4.5–18.5 years of age. These templates provide baseline anatomical models for optical simulations when subject-specific MRI data are unavailable.

For subject-specific head-model construction, anatomical inputs were provided as T1-weighted NIfTI volumes. If acquired in DICOM format, the images were converted to NIfTI prior to import into TLS-Sim. Tissue segmentation was performed using SuperSynth, which generated 1-mm isotropic tissue-label volumes. The native T1-weighted image was registered to the 1-mm MNI152NLin2009cAsym template using SynthMorph, and the resulting spatial transformations were retained for coordinate mapping between subject and standard spaces. All anatomical images and tissue-label volumes were stored in NIfTI format with their associated affine matrices preserved. Voxel indices were mapped to physical coordinates using the corresponding NIfTI affine transformations, thereby maintaining spatial alignment among the anatomical models, light-source locations, simulation outputs, and regions of interest.

For F2F training-data generation, high-quality anatomical segmentations were generated using the Complete Head Anatomy Reconstruction Method (CHARM) within the SimNIBS toolkit [45]. Although CHARM [46] provides a detailed anatomical segmentation containing multiple tissue structures, the native segmentation was subsequently converted into a compact optical-tissue representation comprising six integer classes (five tissue classes plus background/air): background or air, scalp, skull, cerebrospinal fluid, gray matter, and white matter. According to the adopted label convention, native labels 5, 7/8, 3, 2, and 1 were mapped to the scalp, skull, cerebrospinal fluid, gray matter, and white matter classes, respectively. Minor extracranial structures that were not explicitly represented in the compact model, including air cavities and the eyeballs, were assigned to the scalp optical class as a modeling simplification. The skull was retained as a separate optical class and was not merged with the scalp. This conversion preserved the principal anatomical boundaries relevant to intracranial photon propagation while providing a consistent and computationally manageable optical-tissue representation for subsequent Monte Carlo simulation and F2F model training.

### 2.4 Light source placement and direction

TLS-Sim supports pencil-beam, disk, and Gaussian sources for transcranial light-stimulation modeling. Multiple sources can be simulated simultaneously to support multisource and montage-style configurations. Additional light source geometries and parameters can be specified through the simulation configuration file. Light sources can be placed using either subject-specific EEG coordinates or manually specified coordinates. For EEG-guided placement, the selected electrode location, such as Fp1 or C3, is obtained from the subject-specific coordinate table and mapped to the voxel space of the corresponding head model. In the current implementation, the incident direction is determined automatically by orienting the source toward a local cortical target derived from the nearest 5% of gray-matter voxels, with the target defined as the centroid of the three closest gray-matter points along the estimated cortical axis.

For the F2F training and evaluation reported in this study, only pencil-beam sources were used. Source positions were defined using subject-specific EEG 10–20 coordinates, and the incident direction was determined automatically from the local gray-matter geometry beneath each source. Each paired low- and high-photon-count MC simulation used identical source geometry, position, and direction, with only the number of simulated photon packets differing between the two fields.

### 2.5 Tissue optical properties and stimulation parameters

Each optical tissue class was characterized by its absorption coefficient (*μ_a_*, mm⁻¹), scattering coefficient (*μ_s_*, mm⁻¹), anisotropy factor (*g*), and refractive index (*n*). The software includes wavelength-specific tissue optical-property sets for 670, 810, 850, 980, and 1064 nm [47], enabling conventional Monte Carlo simulations at each wavelength. User-configurable stimulation parameters include wavelength, total optical power or irradiance, illuminated area, exposure duration, source position, incident direction, and the number of simulated photon packets. The photon-packet count is specified independently of the physical stimulation dose and primarily determines the statistical noise and computational cost of the simulation. Within TLS-Sim, the Monte Carlo energy-deposition field is normalized to 1 J of total source energy and subsequently rescaled according to the prescribed stimulation parameters. The total delivered optical energy was calculated as⋅ E_delivered_ = I × A × t, where I is the irradiance, A is the effective illuminated area determined by the probe dimensions, and t is the exposure duration. For a circular probe with diameter D, the illuminated area was calculated as 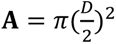. The normalized energy-deposition field was then multiplied by E_delivered_ to estimate the intracranial energy deposition under the specified stimulation conditions.

For the F2F training and evaluation reported in this study, a compact five-tissue optical model was used at 1064 nm. The tissue optical properties were obtained from previously published data and are summarized in Table 1. The absorption and scattering coefficients are expressed in mm⁻¹, whereas the anisotropy factor and refractive index are dimensionless.

**Table 1.** Optical properties assigned to each tissue class in the compact five-tissue head model (1064 nm).

| Tissue | $\mu_a$ (mm<sup>-1</sup>) | $\mu_s$ (mm<sup>-1</sup>) | $g$ | $n$ |
| --- | --- | --- | --- | --- |
| Air / background | 0 | 0 | 1.00 | 1.00 |
| Scalp | 0.0170 | 18.45 | 0.89 | 1.37 |
| Skull | 0.0190 | 14.60 | 0.89 | 1.37 |
| CSF | 0.0144 | 0.09 | 0.89 | 1.37 |
| Gray matter | 0.0530 | 5.90 | 0.91 | 1.37 |
| White matter | 0.1050 | 30.00 | 0.88 | 1.37 |

For the F2F training and evaluation, the representative stimulation configuration used a wavelength of 1064 nm [48–50]. The total delivered optical energy was calculated from the irradiance, effective illuminated area, and exposure duration. Paired Monte Carlo fields were generated using 10⁶ photon packets for the low-photon-count input and 10¹⁰ photon packets for the high-photon-count reference, while all other simulation parameters were held constant.

### 2.6 Monte Carlo simulation and paired-field generation

Photon transport was simulated using MCX through the pmcx Python interface [17]. MCX solves the radiative transport problem through stochastic photon propagation in heterogeneous voxelized media. The simulation configuration specified the tissue-label volume, tissue optical properties, source position and direction, source geometry, photon count, and a single temporal window. The output type was set to deposited energy.

Two simulations with identical anatomy, optical parameters, and source configuration were paired for each sample. For the representative-case comparison, the study used 10^6^ photons for the sparse simulation and 10^10^ photons for the full simulation, respectively. The transformed field and the optical label volume were saved as NIfTI images with the affine matrix of the source image. Each paired training record therefore contained a sparse field and a full field generated from the same subject, source location, and simulation parameters. The sparse field served as the input to the learned model and the full field served as the high-fidelity target.

The released data index comprised 10,328 paired fields drawn from 138 subjects, with 63, 65, or 75 source locations per subject (most subjects contributing 75); each source location provided one sparse input and one full target, so the pairs arose from multiple source locations per subject rather than from repeated targets for a given location. The pairs were partitioned using a subject-level, five-fold scheme in which all fields from a given subject were kept within a single fold, so that the training and test sets of each fold contain disjoint subjects and no subject’s anatomy appears in both.

### 2.7 Flux-to-Flux formulation

The learned acceleration pathway was formulated as a same-domain, F2F transformation. Let *E*_Sparse_ denote the energy-deposition field produced by the low-photon-count MC simulation and *E*_Full_ the corresponding high-photon-count field. The model estimated

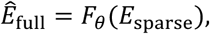

where *F_θ_* was implemented as a latent rectified-flow model. Because *E*_Sparse_ already encoded the effects of the subject’s anatomy, tissue optical parameters, and source configuration through the physical MC simulation, the network mapped between two representations of the same physical quantity rather than translating categorical anatomy into a continuous optical field. This design reduced the cross-modal burden on the learned mapping, but it did not, by itself, establish invariance to demographic group, segmentation method, or optical-property uncertainty; such generalization requires dedicated experiments.

In continuous flow matching [51], a time-dependent vector field *v_θ_*(*x_t_*, *t*) defines an ordinary differential equation

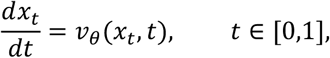

that transports samples between a source distribution and a target distribution. In this paired setting, the sparse-field latent representation supplied the source state and the full-field latent representation supplied the target state. The model was optimized to regress the target velocity along sampled interpolation times [52].

### 2.8 Latent representation and network architecture

Sparse and full log-transformed fields were encoded using a three-dimensional AutoencoderKL derived from the MONAI MAISI implementation [53, 54]. The autoencoder accepted a single input channel and produced four latent channels. Its encoder/decoder channel widths were 64, 128, and 256, with two residual blocks at each resolution and group normalization with 32 groups. The encode–flow–decode structure of this pathway is summarized in Figure 2. The released model checkpoint distributed with the release.

Before encoding, the base-10 log-energy field was clipped to the range [−10,0] and linearly mapped to the autoencoder input range [0,1] as *x*_norm_ = (*x* + 10)/10; the inverse mapping *x* = 10 *x*_norm_ − 10 was applied after decoding to recover the log-energy representation. Encoded latents were multiplied by 0.25 before rectified-flow training and divided by 0.25 before decoding. With the frozen preprocessing configuration used for the reported experiments (an input patch of 160 × 256 × 256 voxels and a spatial downsampling factor of four in each dimension), the resulting latent tensor had four channels and spatial dimensions of 40 × 64 × 64.

The rectified-flow [52] velocity network was a three-dimensional MONAI Diffusion Model U-Net with four input and four output channels. The training script used three resolution levels with channel widths 64, 64, and 128; two residual blocks were used at each level, self-attention was enabled at the deepest level, group normalization used 32 groups, and flash attention was enabled. During training, the sparse-field latent was passed as the source/noise argument of the paired rectified-flow process, and the full-field latent was passed as the target data.

### 2.9 Model training and inference

The repository training script loaded paired latent files by matching filenames ending in simple.pt and full.pt. The full latent was treated as the target and the sparse latent as the source. Training used a batch size of four for 70,000 optimization steps at a learning rate of 3×10⁻⁴, with a maximum gradient norm of 1.0 and exponential-moving-average support inherited from the rectified-flow training framework. Checkpoints were scheduled every 10,000 steps. Inference used the online. The training and inference paths correspond to the solid and dashed arrows in Figure 2.weights, consistent with the loading convention described below.

At inference, the rectified-flow weights were loaded from the online model state of the checkpoint rather than the exponential-moving-average state, following the repository loading convention. The sparse NIfTI field was transformed and encoded by the same autoencoder, multiplied by the 0.25 latent scale, and supplied as the initial/source state to the rectified-flow sampler. The repository default sampling temperature was 1.5. The predicted latent was divided by 0.25, decoded by the autoencoder, clamped to the normalized output range, and mapped back to the log-energy representation. The final prediction was restored to the original NIfTI geometry and saved with the corresponding affine matrix.

### 2.10 Validation design and quality control

The primary comparison was defined among four spatially aligned volumes for each case: the tissue segmentation, the sparse MC field, the full MC reference field, and the fast DL field. Quantitative evaluation was performed in the VAE-decoded [0,1] representation over the whole volume. Peak signal-to-noise ratio (PSNR) was calculated with a fixed maximum value of 1.0. Structural similarity (SSIM) was computed in three dimensions using a Gaussian window (*σ* = 1.5) with stability constants *C*_1_ = (0.01)^2^ and *C*_2_ = (0.03)^2^, as implemented in the released comparison script. Mean absolute error (MAE) and the Pearson correlation coefficient with the full reference were additionally reported. The same metrics were computed for both prediction-versus-full and sparse-versus-full comparisons so that improvement over the physical low-photon baseline could be quantified.

Model validity was additionally assessed using latent-space correlations among the prediction, sparse input, and full target. A usable checkpoint was required to improve prediction-to-target agreement relative to sparse-to-target agreement and to avoid a pass-through solution, in which the prediction reproduces the sparse input (prediction-to-sparse correlation approaching unity) without adding agreement with the full target beyond that already provided by the sparse field. Passing this control therefore requires reporting all three quantities together (prediction-to-target, prediction-to-sparse, and sparse-to-target), because a high prediction-to-target correlation alone does not exclude pass-through when the sparse field is itself already highly correlated with the target. This control was introduced because an archived evaluation of an earlier 70,000-step checkpoint documented pass-through behavior.

Runtime was measured separately for sparse MCX generation, learned inference, and full MCX simulation on the same hardware, with warm-up policy, number of repetitions, and GPU/CPU specifications reported. The spatial error maps were included to complement global image-similarity metrics.

### 2.11 Visualization and dosimetric analysis

TLS-Sim overlaid the simulated or predicted field on the individualized tissue model and supported orthogonal slice views and interactive three-dimensional rendering. Users could adjust the colormap, opacity, display threshold, and slice position. The analysis workflow was designed to quantify energy deposition within selected tissue classes or anatomical regions of interest. Voxel-level inspection combined the energy-deposition field with the optical tissue-label volume in the same NIfTI geometry. Tissue selection enabled users to restrict display and summary calculations to specified labels, while independent thresholds controlled the anatomical overlay and optical field. The software exposed sagittal, coronal, and axial slice controls together with an interactive three-dimensional view, allowing the spatial relationship among the source, skull surface, cortical tissues, and intracranial field to be inspected using a common coordinate system. Analysis inputs and generated summaries were stored separately from the original simulation volume.

For the supplied comparison figures, log-transformed NIfTI fields were loaded into NumPy arrays and visualized using panel-specific procedures. Three-dimensional renderings were used for the volume overview, orthogonal slices were used for structural comparison, and paired absolute-difference maps were used to visualize residual patterns. Any maximum-intensity projection or line-profile analysis is identified separately in the corresponding panel description. The representative comparison used a representative field size of voxels and a log-intensity range of [−10,0].

Depth-dependent attenuation was defined along the incident source direction. Samples were obtained from the scalp entry point toward the brain, and the deposited energy was plotted as a function of depth. Lateral spread was evaluated on planes orthogonal to the incident direction at prespecified depths by quantifying voxels exceeding a defined energy threshold. In the released depth-analysis module the default energy threshold was 10⁻⁶ (decoded units) and the sampling disk radius was 30 voxels, with deposited energy integrated along the sampling path. Group-level comparison functions are included in the software design but have not yet been validated as statistical procedures.

For source-position optimization, the intended workflow would generate a candidate grid on the scalp around the projection of a user-defined intracranial target. A simulation would be evaluated for each candidate position and, when enabled, for multiple source orientations. An ROI objective derived from deposited energy would then be used to rank candidates and return the position and orientation associated with the largest target value. This describes the intended grid-search workflow rather than a validated algorithmic result.

### 2.12 Software and reproducibility

The core dependencies included Python, NumPy, SciPy, nibabel, pandas, pmcx, PyTorch, MONAI, and rectified-flow-pytorch [55]. The runtime environment was verified by direct inspection of the active cluster environment to use Python 3.11.5, PyTorch 2.3.1, MONAI 1.3.2 (with MONAI Generative 0.2.3), NumPy 1.26.4, and pmcx 0.3.5, on an NVIDIA Tesla V100-SXM2-32GB node reporting driver 535.146.02 and a driver-level CUDA compatibility of 12.2 (CUDA runtime wheel 12.1.105).

The code repository contained scripts for MC simulation, latent encoding, rectified-flow training, inference, evaluation, and visualization. Reproducibility of the learned pathway additionally requires the autoencoder and rectified-flow checkpoints, the subject-level split manifest, example input/output volumes, and an executable evaluation command.

## 3. Results

### 3.1 Dataset and evaluation cohort

Under the software-description scope of this article, the Flux-to-Flux pathway is demonstrated on a representative individualized head model rather than evaluated across a large cohort; the training data used to build the released model are summarized here for transparency. The released model was trained on 10,328 paired sparse/full fields from 138 subjects, using a subject-level five-fold split, so that all fields from any given subject fall entirely within one fold and no subject appears in both the training and test sets.

### 3.2 End-to-end TLS-Sim workflow

The complete TLS-Sim workflow was demonstrated from anatomical model construction to simulation, visualization, and quantitative analysis (Figure 1). Starting from a structural T1-weighted image, TLS-Sim generated a five-tissue head model and prepared the anatomical data for subsequent spatial analysis. Stimulation parameters and the target location were then configured within the graphical interface, providing the anatomical and optical inputs required for photon-transport simulation. A continuous end-to-end demonstration of the GUI-based workflow, including model preparation, stimulation configuration, simulation, and subsequent analysis, is provided in Supplementary Video 1.

**Figure 1.**
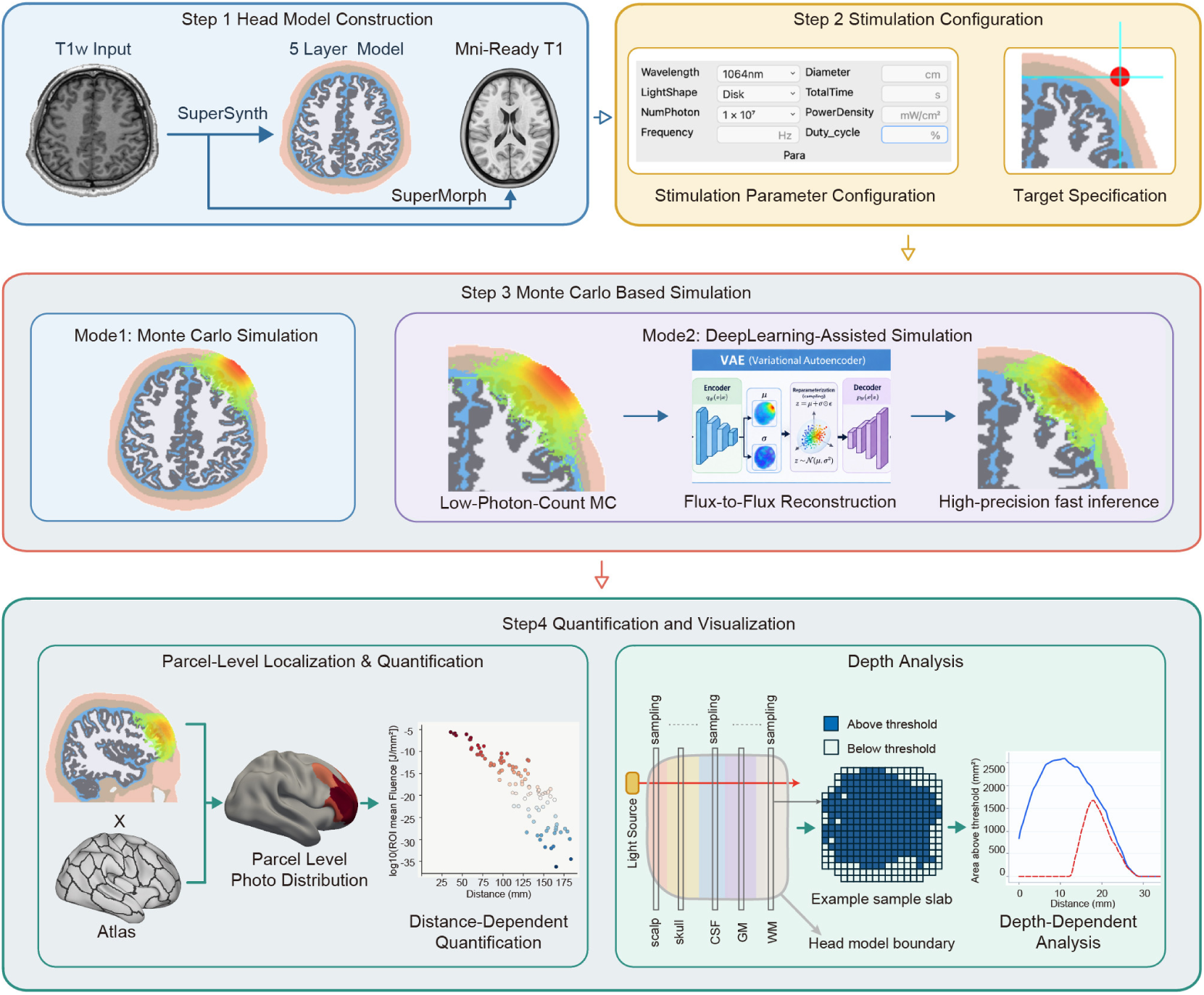
Overview of the TLS-Sim workflow. Step1, head-model construction: structural T1-weighted MRI is segmented into a five-tissue voxelized head model using SuperSynth and spatially registered to MNI space using SynthMorph. Step2, stimulation configuration: wavelength, source geometry, photon-packet count, dose parameters and source placement are configured through the graphical interface. Step3, photon-transport simulation: TLS-Sim provides a conventional GPU-accelerated Monte Carlo pathway and an accelerated F2F pathway. Step4, quantification and visualization: the simulated field is mapped to anatomical parcels for region-level localization and dosimetric quantification, while separate analyses quantify its variation with distance from the source and with depth across tissue layers.

**Figure 2.**
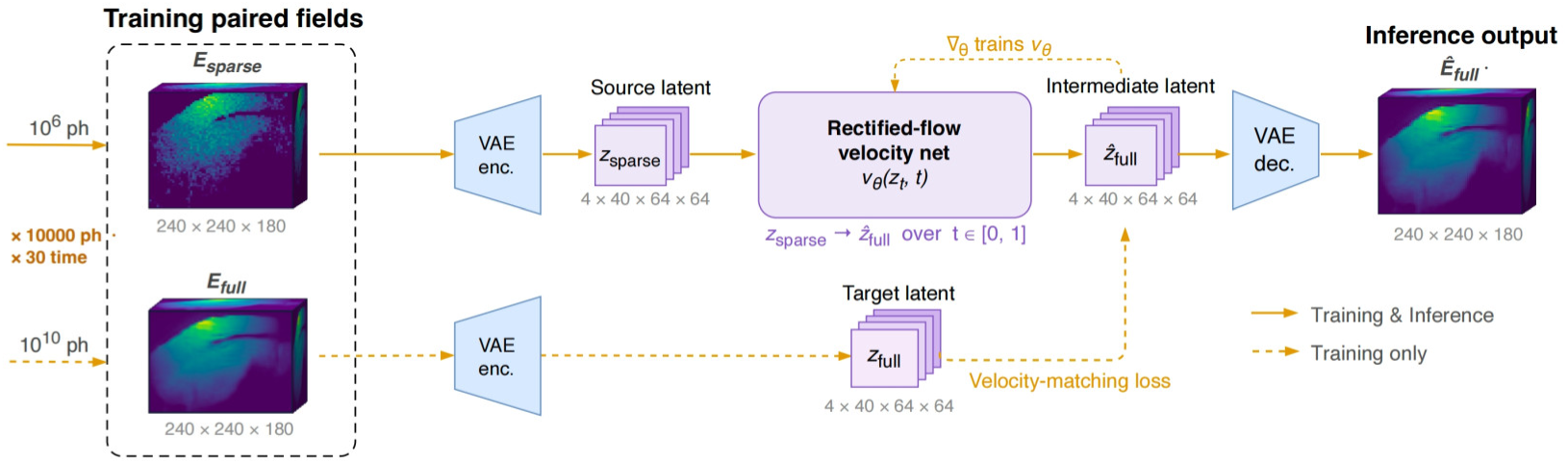
Flux-to-Flux latent rectified-flow refinement. During training (solid and dashed paths), a sparse Monte Carlo field (10⁶ photons) and its paired full-reference field (10¹⁰ photons) are each encoded by a shared, frozen VAE encoder into compact latent tensors (4 × 40 × 64 × 64). A rectified-flow velocity network v_θ_(z_t_, t) transports the sparse-field latent toward the full-field latent along a nearly straight trajectory over t ∈ [0, 1], optimized by a velocity-matching loss in latent space; the VAE decoder reconstructs the predicted full-fidelity field. At inference (solid path) only the sparse field is required, and no full Monte Carlo simulation is run. Fields are shown in the VAE-decoded [0,1] representation for a representative case at 1064 nm.

Following stimulation setup, intracranial light propagation could be estimated using either conventional Monte Carlo simulation or the accelerated F2F pathway. The resulting three-dimensional fluence fields were subsequently integrated with the analysis modules for spatial localization and quantitative characterization. Atlas-based mapping enabled parcel-level identification of the simulated field, while distance- and depth-dependent analyses characterized the spatial attenuation of light relative to the source and across tissue layers. Together, these results demonstrate that TLS-Sim integrates individualized head modeling, stimulation configuration, photon-transport simulation, and dosimetric analysis within a unified end-to-end workflow.

### 3.3 Individualized head-model processing

The anatomical preprocessing pathway accepted a T1-weighted MRI series, converted it to a NIfTI volume, and standardized the working resolution to 1-mm isotropic voxels. The segmentation component produced a multilayer head model containing scalp, skull, CSF, WM, and GM classes, which were subsequently encoded as the discrete optical labels required by MCX. Because the anatomical volume and simulation field retained a common affine transformation, simulation outputs could be overlaid directly on the subject anatomy and interrogated using the same coordinates.

### 3.4 Representative-case comparison across source configurations

The Flux-to-Flux pathway was evaluated on a representative individualized head model for three source configurations, corresponding to electrode positions AF3, AF4, and AF7. For each configuration, a sparse MCX field (10^6^ photons) and a full MCX reference field (10^10^ photons) were generated, and the learned model produced a DL-enhanced prediction from the sparse field. Metrics were computed against the full reference in the VAE-decoded [0,1] representation over the whole volume without a region-of-interest mask. The three fields are shown side by side in Figure 3A, with their differences from the full reference in Figure 3B.

**Figure 3.**
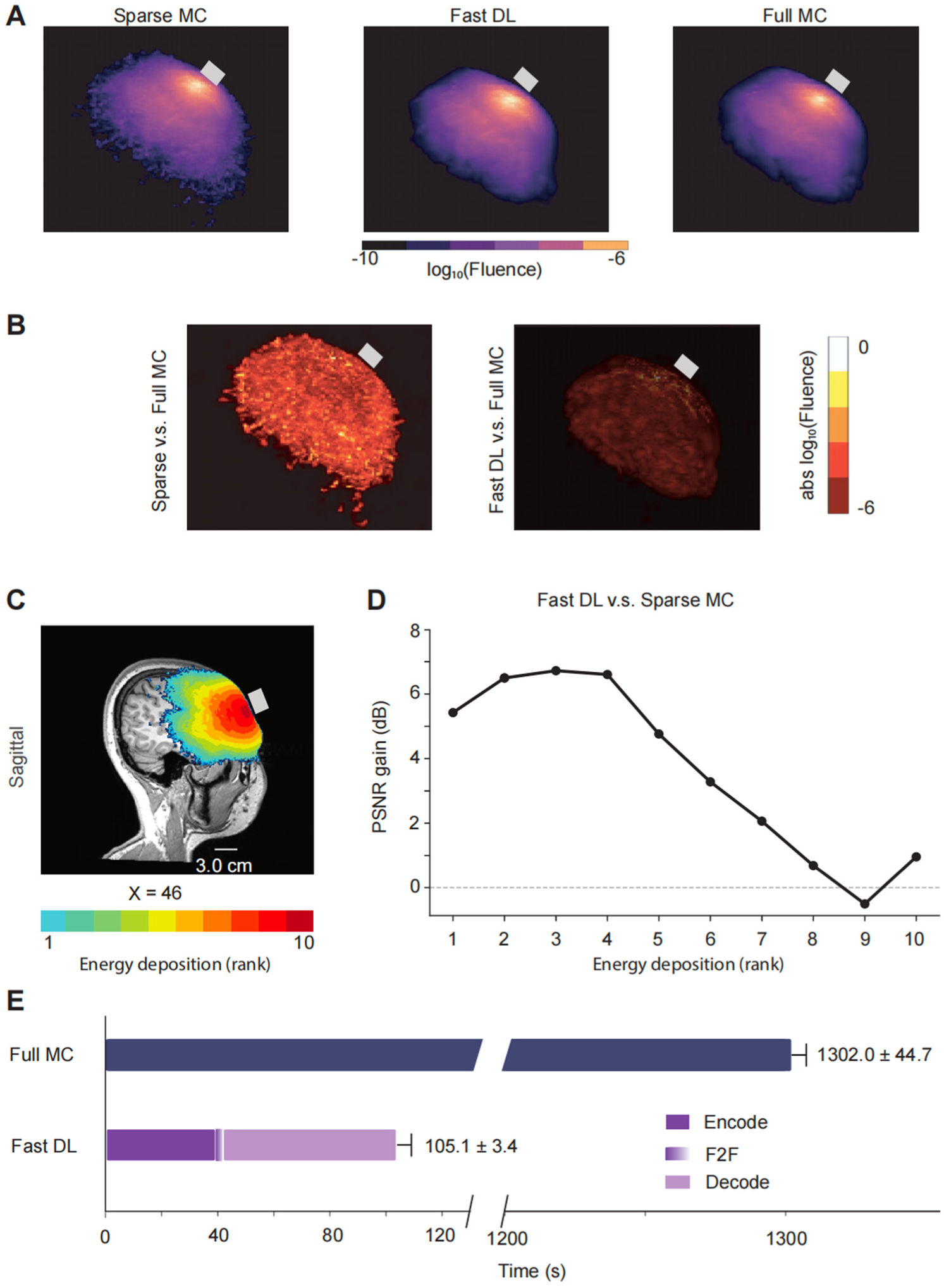
Representative-case evaluation of the Flux-to-Flux pathway (subject IXI012-HH, AF3 configuration, 1064 nm). (A) Volume renderings of the sparse Monte Carlo (Sparse MC), deep-learning-enhanced (Fast DL), and full Monte Carlo (Full MC) log-fluence fields. (B) Absolute log-fluence difference maps relative to the full reference for the sparse field and the DL-enhanced field. (C) Sagittal energy-deposition rank map overlaid on subject anatomy. (D) PSNR gain of the DL-enhanced field over the sparse field as a function of full-reference fluence decile (rank 1 = lowest, 10 = highest); the gain is largest in the lower-fluence boundary deciles. (E) Measured wall-clock runtimes (mean ± s.d., n = 10; NVIDIA RTX 4060, 8 GB) on a broken time axis: full MCX 1302 ± 45 s versus the deep-learning pathway 105 ± 3 s (encode / F2F / decode), an approximately 12-fold reduction.

Across the three source configurations, the learned prediction consistently improved agreement with the full reference relative to the sparse simulation. The primary result is the relative improvement of the prediction over its own physical sparse input: mean PSNR increased by approximately 4.2 dB (range, 3.4–4.9 dB), mean SSIM increased by 0.0057, MAE decreased by approximately 26%, and the Pearson correlation with the full field increased from 0.985 to 0.996. For the AF3 configuration, for example, PSNR increased from 46.01 to 50.92 dB and SSIM from 0.990 to 0.996. Because the sparse and predicted fields share a large low-fluence background and are both expressed in the VAE-decoded [0,1] range, the absolute similarity values are high for both fields; the absolute magnitudes should therefore be read as evaluation-space values. All three configurations derive from the same participant. The per-configuration metrics are summarized in Table 2.

**Table 2.** Representative-case agreement with the full Monte Carlo reference for the sparse simulation and the DL prediction, across three source configurations. PSNR was calculated using a maximum value of 1.0. The reported quantity is the improvement of the DL prediction over the physical sparse input, not the absolute similarity value.

| Configuration | Comparison | PSNR (dB) | SSIM | MAE | Correlation |
| --- | --- | --- | --- | --- | --- |
| AF3 | Sparse vs Full | 46.01 | 0.990 | 0.000908 | 0.984 |
| AF3 | DL vs Full | 50.92 | 0.996 | 0.000634 | 0.996 |
| AF4 | Sparse vs Full | 46.31 | 0.989 | 0.000918 | 0.986 |
| AF4 | DL vs Full | 50.54 | 0.996 | 0.000670 | 0.996 |
| AF7 | Sparse vs Full | 48.20 | 0.993 | 0.000673 | 0.985 |
| AF7 | DL vs Full | 51.64 | 0.997 | 0.000547 | 0.996 |
| Improvement |  | 3.4 to 4.9 | 0.004 to<br>0.007 | −19% to<br>−30% | 0.010 to<br>0.012 |

The visual comparison used the center of the top 1% full-MCX fluence region to define the displayed orthogonal slices (Z = 63, Y = 182, and X = 46 for the axial, coronal, and sagittal views, respectively). The DL-enhanced field reproduced the principal high-fluence structure visible in the full-MCX field across the three views, while the paired absolute-difference maps provided a qualitative visualization of residual error (Figure 3B). The error maps were displayed with a capped color scale and they should be read as qualitative rather than quantitative.

Across the source configurations, the PSNR gain over the sparse input was concentrated at the low-fluence field boundaries, where sparse Monte Carlo shot noise is largest: approximately +6.1 ± 0.1 dB in the lowest-fluence signal voxels versus −0.7 ± 0.9 dB in the highest-fluence core (Figure 3C, D). This spatial pattern is consistent with Monte Carlo variance scaling as the inverse square root of local photon count and indicates that the Flux-to-Flux benefit derives primarily from suppressing low-fluence boundary artifacts rather than from uniform field-wide sharpening. The signal-bearing voxels (full-reference value above a near-zero background threshold) constitute only about 1.3% of the volume, so the whole-volume PSNR gain is weighted heavily by denoising of the near-zero background halo (approximately +5.4 dB there); the improvement inside the energy-deposition field is real but smaller (approximately +3.8 dB) and spatially non-uniform. The whole-volume value should therefore not be read as a uniform, field-wide accuracy improvement.

### 3.5 Pass-through and failure-mode analysis

To confirm that the improvement reflects genuine field refinement rather than a trivial reproduction of the input (a pass-through solution), the predicted field was compared both with its sparse input and with the full reference. The prediction remains strongly correlated with the sparse input, as expected for a same-domain refinement task in which a large low-fluence background is shared (whole-volume prediction-to-sparse correlation ≈ 0.99, and ≈ 0.98 within the high-fluence region). The evidence against pass-through is therefore not a low prediction-to-sparse correlation but the fact that the prediction is measurably closer to the full reference than the sparse input is: in the high-fluence region the prediction-to-full correlation (≈ 0.99) exceeds the sparse-to-full correlation (≈ 0.96), and over the whole volume the prediction improves PSNR by approximately +4 dB and reduces MAE by approximately 26% on average across the three configurations relative to the sparse input (per-configuration MAE reduction 19–30%; Table 2). A high prediction-to-sparse correlation alone does not rule out pass-through, because the sparse field is already highly correlated with the full reference. More importantly, the prediction showed improved agreement with the full reference relative to the sparse input.

### 3.6 Computational performance

Wall-clock runtimes were measured on a single consumer GPU (NVIDIA GeForce RTX 4060, 8 GB) with ten repeated runs per stage (Figure 3E). A full MCX reference simulation (10^10^ photons) required 1302 ± 45 s (approximately 22 min), whereas the deep-learning pathway produced an approximation of the full-reference field in 105 ± 3 s end-to-end, an approximately 12-fold reduction in time to a high-photon-count field. Within the learned pathway, the rectified-flow sampling step itself accounted for only 2.75 ± 0.07 s (16 steps); the VAE encode (39 ± 1 s) and decode (62 ± 2 s) stages dominated inference time. A sparse MCX simulation (10^6^ photons) completed in 1.78 ± 0.06 s, so the learned pathway is not a substitute for a sparse simulation but a means of improving agreement with the full reference from a sparse field at a small fraction of the full-simulation cost.

## 4. Discussion

TLS-Sim unifies the end-to-end workflow of individualized tLS modeling—from anatomical preprocessing and head-model construction to optical-property assignment, source configuration, MC simulation, visualization, and dosimetric analysis—within a single modular Python framework with an integrated graphical interface. By maintaining these steps within a shared, spatially registered volumetric representation, the platform reduces the technical burden associated with data conversion, coordinate transformation, and information transfer among otherwise independent tools, while supporting a more consistent and reproducible modeling workflow. Beyond workflow integration, TLS-Sim incorporates a F2F pathway for accelerated photon-transport estimation. Rather than predicting the intracranial field directly from anatomical data, F2F uses a computationally inexpensive low-photon-count Monte Carlo field, which already encodes the combined effects of individual anatomy, tissue optical properties, and source configuration, as a physics-informed input for reconstructing a high-photon-count reference field. In the representative case, this approach improved agreement with the full field while substantially reducing computational cost. Together, these capabilities help bridge the gap between high-fidelity individualized photon-transport modeling and the computational efficiency required for iterative source optimization and interactive target navigation. The open-source release of TLS-Sim further provides a shared and extensible foundation for independent validation, method comparison, and continued community development.

In this study, we illustrate how the framework integrates the computational workflow and evaluates the accelerated pathway. First, we show that preprocessing, source and head modeling, simulation, and dosimetry share a common volumetric workflow; Second, we set out the Flux-to-Flux latent rectified-flow pathway that produces the accelerated field estimate; and Figure 3 summarizes the representative-case evaluation, from field and difference maps through the fluence-stratified gain to the measured runtimes. TLS-Sim maintains preprocessing outputs, head models, source definitions, simulated fields, and dosimetric results within a shared volumetric coordinate system, allowing intermediate products to be inspected and reused. The graphical interface was implemented as a front end to the underlying computational modules, allowing users to configure and execute simulations without directly modifying backend files. This separation improves usability, workflow transparency, and software maintainability while preserving a consistent numerical backend across different operations.

The F2F formulation reframes the task as a learned denoising problem in the Monte Carlo simulation domain. Unlike anatomy-to-field emulators, F2F does not predict photon transport directly from anatomical images. Instead, it uses a computationally inexpensive low-photon-count MCX field as a physics-informed input. A sparse MCX field already reflects the interaction among individual anatomy, assigned optical properties, and the selected source configuration. Using that field as the learned input therefore preserves a physics-based initialization and reframes the neural-network task as the refinement of a noisy, under-sampled field rather than its synthesis from anatomy. This within-domain formulation is conceptually appealing because it reduces the learning burden while anchoring the reconstruction to a physics-based Monte Carlo estimate, but it does not by itself guarantee robustness across populations, segmentation methods, wavelengths, or tissue-property uncertainty; establishing such robustness requires explicit out-of-distribution experiments.

The representative comparison across three source configurations provides a useful proof-of-workflow observation: in the VAE-decoded representation, the DL output improved over the sparse MCX baseline in PSNR, SSIM, MAE, and correlation against the paired full-MCX field, with a mean PSNR gain of approximately 4 dB. The accompanying orthogonal slices show that the high-fluence pattern is preserved. Together these results provide proof-of-concept evidence that F2F can refine an inexpensive physical estimate into one that agrees more closely with the full simulation. However, the findings should be interpreted cautiously because the evaluation was limited to a single individualized head model tested across multiple source placements, and global image metrics may be strongly influenced by the large number of low-fluence background voxels. The most informative result is therefore the improvement relative to the low-photon-count MCX baseline rather than the absolute similarity to the reference field.

A fluence-stratified analysis clarifies the mechanism behind this improvement. The refinement behaves as a variance-aware denoiser: its benefit is concentrated at low-fluence field boundaries, where the sparse simulation’s shot noise is largest, and is essentially neutral at the high-fluence core that the sparse field already resolves well. This is the behavior expected when Monte Carlo variance scales as the inverse square root of local photon count, and it recasts F2F not merely as a full-simulation accelerator but as a learned suppressor of sparse-simulation boundary artifacts. The same analysis is a reminder that the whole-volume metric is background-weighted and that the high-fluence core is improved least; reporting fluence-stratified or signal-restricted metrics is therefore both more informative and more defensible than the whole-volume figure alone.

The measured end-to-end acceleration warrants a precise interpretation. On identical consumer hardware, the complete F2F reconstruction pathway was approximately 12 times faster than the high-photon-count MCX simulation. The reported runtime included VAE encoding, Flow Matching inference, and decoding, rather than reporting the latent sampling time alone. Flow Matching inference required less than 3 s, whereas the encoding and decoding stages accounted for most of the inference time, identifying the autoencoder as the principal target for further optimization. Because the low-photon-count MCX simulation itself completed in under 2 s, F2F should be viewed not as a replacement for Monte Carlo simulation, but as a pathway from a rapid physical estimate toward high-photon-count reference quality without incurring the full simulation cost. The source-position optimization concept could extend TLS-Sim from forward simulation to treatment-parameter search. This acceleration is particularly relevant to iterative source-position optimization and interactive target navigation. Conventional high-photon-count simulation makes repeated evaluation of large numbers of candidate source positions and orientations computationally expensive. Integrating F2F with ROI-based optimization may allow candidate stimulation configurations to be compared within a more practical time frame. However, this application remains prospective and will require clearly defined optimization objectives, source-placement constraints, baseline comparisons, and sensitivity analyses for grid resolution and optical-property uncertainty before treatment-planning performance can be established.

Several limitations remain. First, the F2F pathway was evaluated only as a proof of concept using a single individualized head model, and its generalizability across subjects and anatomical variations remains to be established. Second, the framework uses a compact five-tissue optical model in which minor extracerebral structures are approximated by the scalp optical class. Consequently, segmentation errors, tissue simplifications, and uncertainties in the assigned optical properties affect both the low- and high-photon-count simulations. Moreover, the high-photon-count Monte Carlo field remains a numerical reference rather than an in vivo ground truth. Therefore, the predicted intracranial dose should be interpreted as a model-based estimate, and agreement with the reference field does not establish absolute dosimetric accuracy, biological efficacy, or clinical validity. Third, the voxel-based representation may not fully capture thin tissues, curved interfaces, or subvoxel anatomical features. Finally, the current Flux-to-Flux model has been trained and validated only at 1064 nm. Additional wavelengths, source profiles, and advanced functions, including source optimization, obstruction compensation, and group-level analysis, require dedicated validation before their performance can be generalized. These capabilities will be incorporated into future releases as the corresponding methodological frameworks are established and published.

Priorities for future work are therefore to test generalization to unseen subjects and to a broader range of anatomies, segmentation pipelines, optical-property assumptions, wavelengths, and source configurations. Comparisons with mesh-based simulations and experimental measurements will also be needed to confirm dosimetric accuracy beyond agreement with a numerical reference, while prospective studies should determine whether faster simulation meaningfully influences stimulation-planning decisions. Most directly, the acceleration demonstrated here provides an enabling step toward three capabilities that are difficult to achieve with conventional computationally intensive simulations: rapid iterative source-position optimization, obstruction compensation, and interactive target navigation. Building these capabilities on top of the Flux-to-Flux pathway represents a natural next stage, with the broader goal of moving individualized photon-transport simulation from offline batch computation toward near-interactive planning within an open and extensible TLS-Sim platform.

## 5. Conclusion

TLS-Sim provides a modular framework for individualized transcranial light-stimulation simulation, visualization, and dosimetric analysis. Its Flux-to-Flux pathway uses a sparse Monte Carlo field as a physics-informed starting point for estimating a high-photon-count field and is demonstrated here on a representative case. On this basis, TLS-Sim offers a practical research platform for developing and comparing individualized tLS protocols. The open-source release of the toolkit, trained model, and representative data enables users to reproduce the reported results and evaluate the pathway using their own datasets. Broader subject-independent validation of reconstruction accuracy and computational efficiency remains necessary.

## Data and Code Availability

TLS-Sim is released as an open-source Python toolkit so that the workflow and the representative results reported here can be independently reproduced and extended. The trained rectified-flow checkpoint and the variational-autoencoder weights used for all reported results, together with the per-configuration sparse, DL, and full NIfTI volumes and the machine-readable benchmark record for the representative subject, are archived with the release.

## Ethics Statement

This study used structural MRI data from the publicly available IXI dataset (Information eXtraction from Images; https://brain-development.org/ixi-dataset/), which is released for research use under a Creative Commons license. All images in this dataset are de-identified at source. No new human or animal data were acquired for this work, and no additional institutional review board approval was required for the secondary analysis of these openly licensed, anonymized images.

## CRediT authorship contribution statement

Keyao Zhang, Hai Jia and Zhilin Li contributed equally.

- **Keyao Zhang:** Conceptualization, Software (toolkit and simulation pipeline), Methodology, Visualization, Writing – original draft.
- **Hai Jia:** Software, Methodology, Formal analysis, Data curation, Writing – original draft.
- **Zhilin Li:** Supervision, Writing – review & editing.
- **Yushuai Zhang:** Investigation, Data curation, Visualization.
- **Xiaoli Li:** Supervision, Resources, Funding acquisition.
- **Chenguang Zhao:** Conceptualization, Supervision, Project administration, Writing – review & editing (corresponding author).

## Declaration of generative AI and AI-assisted technologies in the manuscript preparation process

During the preparation of this work, the authors used ChatGPT (OpenAI) to improve the language, clarity, and readability of the manuscript. After using this tool, the authors reviewed and edited the content as needed and take full responsibility for the content of the publication.

